# The Genetic and Evolutionary Landscape of Pentanucleotide Tandem Repeats in Human Genomes

**DOI:** 10.1101/2025.05.27.656002

**Authors:** Isaac R. L. Xu, David Pellerin, Liedewei Van De Vondel, Matt C. Danzi, Stephan Züchner

## Abstract

Tandem repeats are a class of genetic variation characterized by repetitions of DNA motifs of diverse sequences and lengths. Pentameric motifs, in particular, do not align with the protein-coding reading frames, often confining them to the less extensively studied non-coding portion of the genome. Intronic pentanucleotide tandem repeats are associated with 11 neurological disorders. Their pathogenic genotypes are primarily characterized by large length expansion and, unlike most repeat expansion disorders, have required a deviation from the reference motif. The population-level variability of pentameric repeats remains poorly understood due to technical limitations of short-read sequencing technologies. To address this knowledge gap, we genotyped 28,446 pentanucleotide repeat loci in 1,027 long-read PacBio HiFi samples from self-identified Black and African American participants in the All of Us Research Program. We developed new algorithms for tandem repeat decomposition, characterization, and visualization to facilitate this analysis. Our findings reveal extensive genome-wide heterogeneity in repeat length and sequence composition. Alleles with DNA sequences containing segments of distinct motifs were observed in 15% of loci. Additionally, 8% of loci exhibited multimodal repeat length distributions, in which distinct sequence compositions were often associated with distinct length ranges. Repeat loci were highly enriched in proximity to transposable elements, including 68% mapping to Alu elements, a retrotransposon specific to primates. Comparative analysis of the reference sequences from 30 species, including 27 primates, suggests that most pentanucleotide repeats fully emerged in a common ancestor of humans and other ape species, or even within more closely related hominin lineages. As an example, we describe a highly polymorphic and recently evolved repeat locus in the *ABCA1* gene, a major regulator of cellular cholesterol and phospholipid homeostasis. This study provides novel and comprehensive insights into the evolutionary formation of pentamer tandem repeat loci and their extensive variability across human genomes.

## INTRODUCTION

Tandem repeats constitute ∼8% of the genome and contribute to an expanding number of diseases mainly implicated in neurological phenotypes[1, 2]. Due to their repetitive nature, the mutation rate of tandem repeats is much higher than the standard mutation rate, making the among the most polymorphic forms of variation in the human genome [3]. Despite their abundance and clinical relevance, tandem repeats remain one of the most enigmatic forms of genetic variation, owing largely to challenges in their detection and analysis.

Traditional genomic techniques have significant limitations when analyzing tandem repeats. Methods like Sanger sequencing often fail with large tandem repeats and require precise specifications for oligonucleotide primer design and amplification protocols. Southern blotting and PCR-based gel electrophoresis can approximate repeat length but lack information on sequence composition and are low-throughput. Over the past decade, short-read sequencing technologies have dominated genetic studies. While effective for shorter tandem repeats, the limited read-length prevents accurate characterization of repetitive sequences larger than ∼150 base pairs. Further, short-read genotyping software often requires prior catalogued specification of repeat motif and structure [4], limiting its ability to genotype novel motifs and impure repeat alleles in large genome-wide screens. Tools such as ExpansionHunter Denovo and STRling, developed for non-catalog-based repeat detection in short-reads, may indicate the presence of an expanded allele, yet cannot accurately resolve its size or sequence [5, 6].

The current knowledge of tandem repeat variation in the population has been informed by large-scale sequencing experiments [7–11]. However, most population-level studies have relied on short-read sequencing technologies. Further, most previous studies often simplify a tandem repeat sequence as a motif unit and count to facilitate large scale genotyping [7, 8]. This approach often uses the reference genome sequence as a model for repeat structure, which may not capture the full range of variability, such as interruptions or alternate motifs. This limitation is particularly significant considering the sequence content of a tandem repeat can profoundly impact key clinical features such disease onset, penetrance, and severity [12–19]. These challenges highlight the need for more nuanced methods to analyze tandem repeats at the sequence level, moving beyond simple metrics to uncover their role in health and disease.

Intronic pentanucleotide tandem repeats (PNTR), currently associated with 11 repeat expansion disorders, exemplify the limitations of current methodologies. These disorders are driven by expansions containing rare motifs that are absent from the reference genome[12, 13, 20]. The known disorders include Familial Adult Myoclonic Epilepsy (FAME) Types 1, 2, 3, 4, 6, 7, and 8; Spinocerebellar Ataxia Types 10, 31, and 37; and Cerebellar Ataxia with Neuropathy and Vestibular Areflexia Syndrome (CANVAS). All but one of these loci map to an Alu element, a retrotransposon unique to primates. Pathogenic alleles often exhibit complex, chimeric repeat sequences, in which the disease-causing motif is embedded within a larger repeat structure that includes reference or other motifs. For example, at the seven FAME loci, patient alleles may contain configurations such as (TTTTA)_n_(TTTCA)_m_ or nested (TTTTA)_n_(TTTCA)_m_(TTTTA)_k_ or alternative (TTTTA)_n_(TTTCA)_m_(TTTTA)_k_(TTTCA)_j_, where TTTTA is the reference motif [21]. The proportions of pathogenic and reference motif units can vary significantly between patients, with the pathogenic motif accounting anywhere from over 90% to as little as 1% of the total repeat sequence in reported cases [12, 22, 23]. Benign expansions containing reference or other motifs have been observed at these loci, sometimes in over 5% of control subjects, creating an additional complication in distinguishing benign from pathogenic expansions [24, 25]. As it stands, standard short-read genetic techniques are inadequate for characterizing these 11 loci and are unlikely to support the discovery of novel loci. Indeed, identifying most of these genetic disorders relied on well-defined linkage regions, multiple affected pedigrees, and manual inspection of long-read sequencing results [24, 26–31].

The advent of long-read sequencing has the potential to revolutionize the study of tandem repeats. With read lengths exceeding 10kb, long-read sequencing can, in most instances, fully span a repetitive allele, enabling base-pair resolution of the tandem repeat sequence. This study conducted a comprehensive genome-wide analysis of 1,027 PacBio HiFi genomes from the All of Us research program, focusing on pentanucleotide repeats—a class of tandem repeats ideally suited for sequence-level analysis due to critical role of alternate repeat motifs in driving pathogenicity. To enable this work, we developed innovative approaches and metrics to segment, characterize, and visualize tandem repeats, facilitating detailed analyses at base-pair resolution and providing a robust framework for investigating these loci. By leveraging the high accuracy and capabilities of modern long-read sequencing technologies, this study elucidates population-level variation, identifies features distinctive of pathogenic loci, and provides novel insights into the genetic architecture of a currently enigmatic form of variation.

## RESULTS

### Pentanucleotide repeats (PNTR) are enriched in introns and preferentially map to *Alu* elements

A total of 28,446 PNTR loci from 1,027 long-read PacBio HiFi samples were identified and genotyped (see Methods). Among these loci, 56% were located within introns, 43% in intergenic regions, and 1% were in the untranslated regions of genes. Over 58 million alleles were decomposed to enable a detailed analysis of their length and content (see Methods). On average, the human genome in this cohort contained 393.22 base pairs of PNTR per megabase, with chromosome arms 19p and 16p exhibiting the highest density of genic repeats (Figure 1A). In contrast, repeat densities were lower relative to chromosome length on the acrocentric chromosomes (13, 14, 15, 21, and 22) as well as on 18q, 9q, 1q, and Xq. Repeat density calculations were confined to non-centromeric and non-telomeric regions. The distributions of repeat lengths between genic and intergenic loci were largely similar, with mean lengths of 40.1 and 42.7 bases (SD = 6.1 and 6.3), and major allele frequencies of 0.713 and 0.689, respectively (Figure 1B).

**Figure 1:**
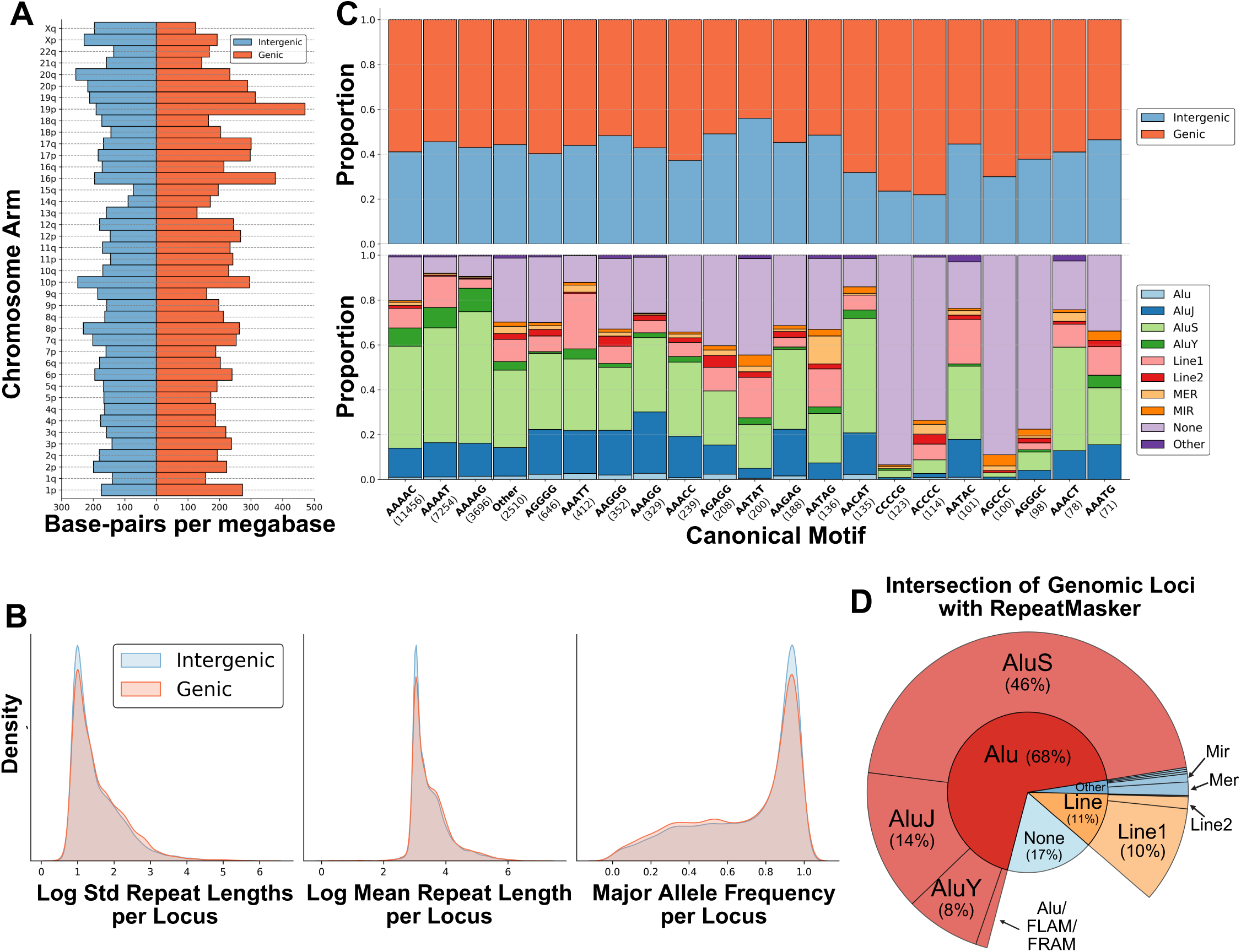
Distribution of pentanucleotide tandem repeats across the genome. A) Diverging bar plot of pentanucleotide tandem repeat density across chromosome arms. The x-axis represents the number of base pairs per mega base occupied by tandem repeats, while the y-axis denotes chromosome arms. Tandem repeats located in genic regions are shown in red, and intergenic repeats are shown in blue. B) Kernel density estimation of repeat locus characteristics. Density distributions are shown for the log standard deviation of repeat lengths (left), log mean repeat length per locus (middle), and major allele frequency per locus (right). Genic tandem repeats are represented in red, and intergenic tandem repeats are in blue, with overlapping areas blending. C) Stacked barplot of the distribution of tandem repeats across different canonical motifs. The top panel shows the proportion of genic (red) and intergenic (blue) repeats for the 20 most frequent motifs, with motif frequency indicated in parentheses on the x-axis. Motifs outside the top 20 are grouped under “Other”. The bottom panel presents the overlap of these motifs with RepeatMasker-annotated elements. Loci with no repeatMasker annotation was marked as “None”. D) Sunburst chart depicting the RepeatMasker annotation of 28,446 genomic pentanucleotide repeat loci. The inner ring represents broader categories, while the outer ring provides finer classifications.

The canonical motif was defined as the motif with the largest base-pair contribution within a repeat sequence (primary motif) across the majority of alleles at a locus. Among all loci, 93 of 102 possible canonical pentanucleotide motifs were identified when simplifying for frame and strand orientation (see Methods). The canonical motifs AAAAC, AAAAT, and AAAAG were the most common, collectively accounting for 78.77% of all loci (Figure 1C). Including these three, only four other canonical motifs were present in over 1% of loci, while 50 canonical motifs were identified in 10 or fewer loci. Additionally, 2.9% and 0.9% of loci contained tetranucleotide and hexanucleotide motifs as the canonical motif, respectively. While most canonical motifs were similarly distributed in genic vs intergenic loci, CG-rich motifs such as CGGGG, ACCCC, AGGGC, and AGCCC were more frequently observed in genic loci (Figure 1C).

Of the eleven currently known disease-associated pentanucleotide repeat loci, ten map to Alu elements (Table 1). To place this in context, the coordinates of all loci were intersected with the RepeatMasker hg38 database. Over 80% of loci overlapped a transposable element (Figure 1D), with 68.4% mapping specifically to an Alu element. The wildtype motifs observed at these pathogenic loci such as AAAAG, AAAAT, and AACAT exhibit a genome-wide preferential association with Alu elements, with 85.2%, 76.7%, and 75.6% of loci containing these canonical motifs intersecting an Alu element, respectively (Figure 1C). Loci that did not overlap with Alu elements were enriched for CG-rich motifs. Alu elements contain a central polyadenine region and a distal polyadenine tail at their 3′ end; among loci that map to Alu elements, approximately 96% fall within the polyadenine tail, while 4% localize to the central polyadenine region.

**Table 1:** Summary of disease-associated intronic pentanucleotide loci (N = 11). *Pathogenicity of the AAAGG repeat in *RFC1* has been observed when the expansion exceeds 500 repeat units and is present in a compound heterozygous state with an AAGGG expansion.

| Gene | Wildtype Motif | Pathogenic Motif | Year Published | Disease | RepeatMasker Overlap |
| --- | --- | --- | --- | --- | --- |
| *ATXN10* | AATAG | AATGG | 2000 | SCA10 | AluSx1 |
| *BEAN1* | AAAAT | AATGG | 2009 | SCA31 | AluSx |
| *DAB1* | AAAAT | AAATG | 2017 | SCA37 | AluJb |
| *SAMD12* | AAAAT | AAATG | 2018 | FAME1 | AluSq2 |
|  |  | AAATC |  |  |  |
| *TNRC6A* | AAAAT | AAATG | 2018 | FAME6 | AluSx1 |
| *RAPGEF2* | AAAAT | AAATG | 2018 | FAME7 | None |
| *RFC1* | AAAAG | AAGGG | 2019 | CANVAS | AluSx3 |
|  |  | ACAGG |  |  |  |
|  |  | AGGGC |  |  |  |
|  |  | AAAGG* |  |  |  |
| *STARD7* | AAAAT | AAATG | 2019 | FAME2 | AluSx1 |
| *MARCHF6* | AAAAT | AAATG | 2019 | FAME3 | AluSx |
| *YEATS2* | AAAAT | AAATG | 2019 | FAME4 | AluSx |
| *RAI1* | AAAAT | AAATG | 2024 | FAME8 | AluJb |

In summary, loci harboring PNTR are a genome-wide ubiquity and enriched in genes. They predominantly map to *Alu* elements, making them a potential source of evolutionary recent variation, unique to certain primate genomes.

### Length heterogeneity of pentanucleotide repeats

Among the 28,446 loci analyzed, varying degrees of allele length heterogeneity were observed. Most loci contained minimal degrees of polymorphism. Specifically, the majority of loci (14,764; 51.9%) exhibited a difference of 10 or fewer base pairs between their maximum and median allele lengths, while 26,207 loci (92.1%) had differences of 50 or fewer base pairs. By contrast, only 727 loci (2.56%) exceeded 150 base pairs in this metric.

A documented phenomenon regarding PNTR disease-associated loci is the presence of benign expansions of the reference motif in the general population [24, 25]. To maintain consistency with Cen et al. regarding their measurement of *SAMD12* AAAAT expansions, we defined an “expansion” as ≥400 base pairs (∼80 pentameric units). In this cohort, 10 of the 11 disease loci contained expansions of either the canonical motif or an alternate motif (Table 2). In most cases, these were observed in fewer than 1-2% of alleles, except for the *RFC1* locus, where 15.64% of alleles had an AAAAG expansion, and 1.37% exhibited an expansion with a different primary motif. The *SAMD12*-FAME1 repeat locus was the only disease-associated locus without an expansion of either the canonical or an alternate motif in this cohort, despite being reported in 3.4-5.9% of individuals in other studies. In a separate cohort of 465 Oxford Nanopore Technology (ONT) genomes that included more ancestries (see Methods), we observed that *SAMD12* expansions were more commonly found in east and south Asian populations (Supplementary Figure 1).

**Table 2:**
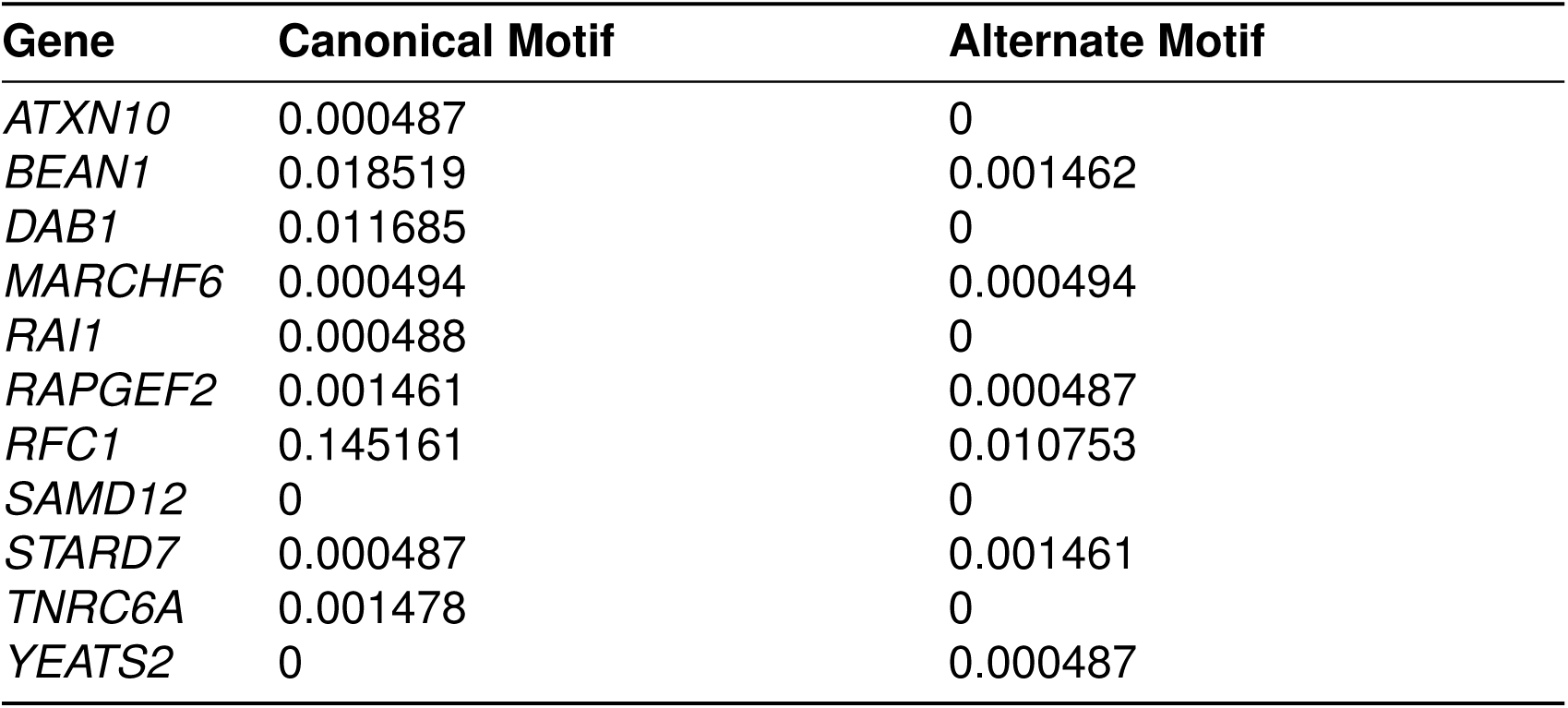
Expansion frequency of disease-associated pentanucleotide loci in a self-identified Black and African American cohort (N = 1,027). Frequencies reflect the proportion of alleles exceeding 400 bp in total repeat length, where the primary motif — defined as the motif contributing the most base pairs to the allele — matched either the canonical (wildtype) or an alternate motif.

To further quantify the presence and extent of expansions beyond known pathogenic loci—and as a proxy for allelic instability—an expansion score was calculated as follows:

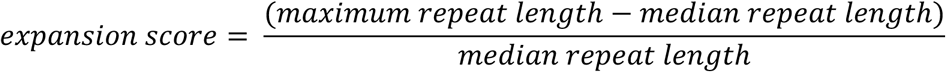

The median expansion score among all loci was 0.4. However, 9 of the 11 disease-associated loci had expansion scores above the 99^th^ percentile score of 8.75 (Figure 2A). When grouped by canonical motif, most motifs had very few loci with elevated expansion scores (Figure 2B). Yet, loci with AAAAT as the canonical motif were highly represented among the highest expansion scores. Consistently, AAAAT is the canonical motif in 9 of 11 disease loci. The largest expansion score identified in this study corresponds to a locus in intron 12 of the *MORN1* gene and had a rare canonical motif of AGCCC. Despite a median length of only 3 units, some alleles extended to nearly 4,000 base pairs at this locus. Unlike *SAMD12*, expanded alleles in *MORN1* were more often observed in African genomes (Supplementary Figure 2).

**Figure 2:**
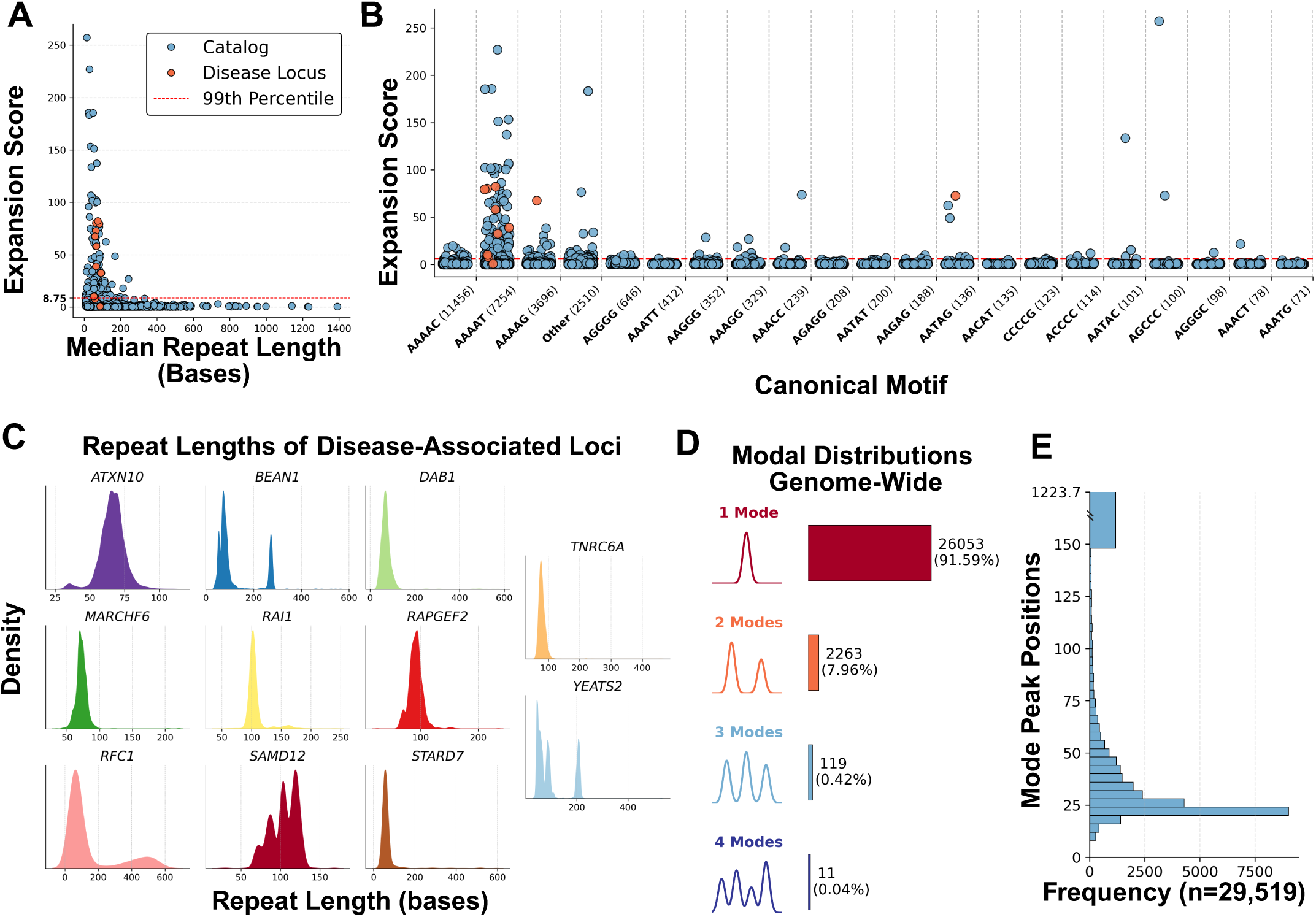
Length heterogeneity of pentanucleotide tandem repeats genome-wide. A) Scatterplot illustrating the expansion score, defined as 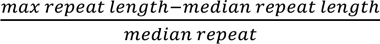 for 28,446 pentanucleotide repeat loci. Disease-associated loci (N=11) are highlighted in red, while all other cataloged loci are shown in blue. The red dashed line marks the 99th percentile expansion score threshold (8.75). The x-axis represents the median repeat length at each locus, and the y-axis denotes the expansion score. B) Scatterplot of expansion scores across different canonical pentanucleotide repeat motifs. Each point represents an individual repeat locus, with disease-associated loci highlighted in red. The 99th percentile expansion score is denoted by the red dashed line. Canonical motifs are displayed on the x-axis, ordered by frequency (shown in parentheses). C) Kernel density estimation (KDE) plots of repeat length distributions for 11 disease-associated loci. Each subplot represents a different gene, labeled above the corresponding plot. The x-axis denotes repeat length (in base pairs) for all repeats at the locus, restricted to <600 bp. The y-axis represents the density of the distribution. D) Distribution of multimodal repeat length patterns genome-wide. The left side illustrates example density plots for unimodal, bimodal, trimodal, and tetramodal distributions. The corresponding colored bars represent the proportion of loci exhibiting each type of distribution. The total number of loci and their respective percentages are annotated on the right. E) Horizontal barplot of modal repeat length peaks (n = 29,519) detected across 28,446 loci. The x-axis represents the frequency of loci with a given mode, while the y-axis indicates the repeat length in base pairs. All modal peaks greater than 150 bp, up to the maximum of 1223.7 bp, were grouped into a single bin.

Loci with large median lengths (>150 base pairs) generally exhibited smaller differences between median and maximum lengths and were often composed of AG-rich motifs. The motifs AGGGG and AAGGG tended to present in impure clusters of variation that contained multiple short motif segments. In contrast, AAAGG was capable of forming longer, pure segments across multiple loci. For instance, a repeat locus in intron 5 of *MID2* displayed high polymorphism, with a major allele consisting of 44 uninterrupted copies of the AAAGG motif.

### Identifying multimodal distributions of repeat length

Visualization of decomposed repeat lengths at the 11 disease-associated loci revealed multimodal distributions in multiple instances, including a bimodal distribution in *RFC1* and *BEAN1* and trimodal distributions in *YEATS2* and *SAMD12* (Figure 2C). To quantify the presence of multiple repeat length modes at a locus, Kernel Density Estimation (KDE) and peak detection were applied (see Methods). Across all loci, 24,772 loci (91.59%) exhibited a unimodal distribution, whereas 2,189 loci (7.96%) had a bimodal distribution (Figure 2D). Tri- and tetra-modal distributions were much rarer, occurring in fewer than 1% of loci.

The positions of these modal peaks show that the majority of PNTRs cluster at relatively small lengths, with a median of 27 and a mean of 43.8 base pairs (Figure 2E). The maximal repeat length peak observed in this study was 1223.7 base pairs. At disease-associated loci, peaks ranged from 35.4 to 271.7 with a median of 72 and a mean of 91.5 base pairs (Supplementary Figure 3).

### Sequence heterogeneity at pentanucleotide repeat loci

#### Deriving common sequence haplotypes at a locus

Sequence analysis of PNTR demonstrated that distinct motif and interruption patterns (sequence configurations) dominate certain length ranges and are the source of multimodal distributions (Figure 3A, 3D). To derive common sequence configurations at a locus, alleles with matching order of repeat motifs and interruptions were grouped (see Methods). We chose two loci, *YEATS2*-FAME4 and *SAMD12*-FAME1, to illustrate the effect sequence composition can have on interpretation of repeat length (Figure 3B,3E).

**Figure 3:**
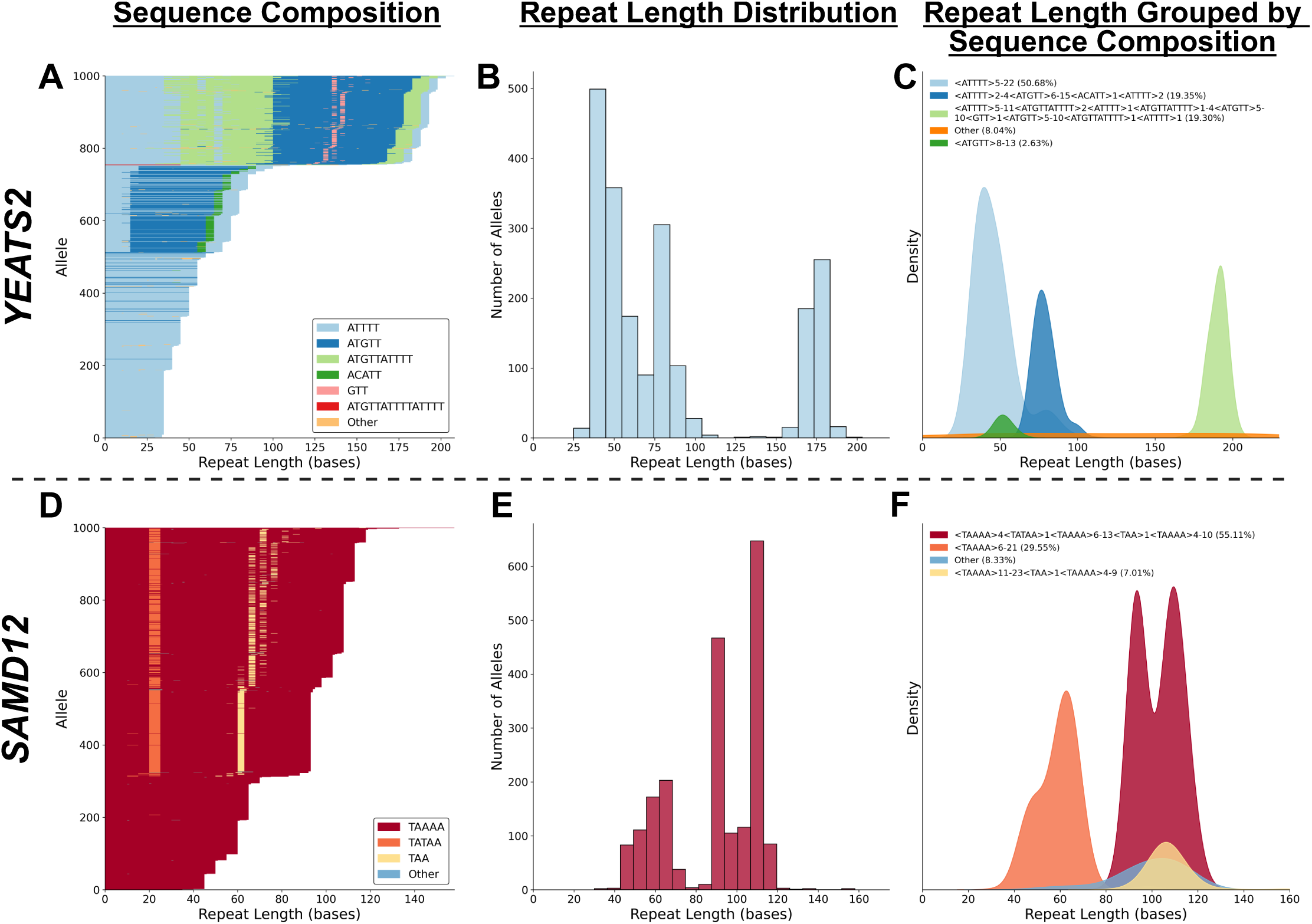
Distinct sequence compositions driving multimodal distributions at a locus. A, D) Waterfall plots displaying the allelic composition of 1,000 randomly selected alleles at the *YEATS2-*FAME4 (A) and *SAMD12-*FAME1 (D) repeat loci. Each row represents an individual allele, with colored bars corresponding to specific repeat motifs or interruptions as indicated in the legend. The x-axis represents the repeat length in base pairs. B, E) Histograms of decomposed repeat lengths at the *YEATS2*-FAME4 (B, n=2,052) and *SAMD12*-FAME1 (E, n=2,054) loci. The x-axis represents repeat length in base pairs, while the y-axis indicates the number of alleles. C, F) Tide plots (grouped density plots) of repeat length distributions at the *YEATS2-*FAME4 (C) and *SAMD12-* FAME1 (F) loci, stratified by distinct repeat sequence configurations. Colors correspond to unique repeat structures as specified in the legend, where structures are represented as <Motif>min,max. The min and max values denote the minimum and maximum observed motif copy numbers for alleles with that structure. If min=max, only a single value is shown. The x-axis represents repeat length in base pairs.

At the *YEATS2*-FAME4 locus, 50 percent of alleles contained pure ATTTT repeats between 5 and 22 units (median of 9), consistent with the hg38 reference. However, three additional configurations were identified (Figure 3C). One structure, found in approximately 19 percent of chromosomes, was tightly clustered between 170 and 200 base pairs (interquartile range of 5) and contained ATTTT, ATGTT, and ATTTTATGTTT motifs. These distinct sequence structures each occupied narrower and partially non-overlapping length ranges, indicating that repeat size distributions are strongly influenced by the underlying sequence composition. In the 465 Oxford Nanopore genomes from the 1000 Genomes Project, which included greater ancestral diversity, the frequency of these sequence configurations differed by superpopulation (Supplementary Figure 4).

At the *SAMD12*-FAME1 locus, three major peaks were observed at 63.3, 93.2, and 108.7 base pairs, corresponding to three dominant sequence configurations (Figure 3E). Only 7 percent of alleles matched the hg38 reference structure. A closely related configuration, differing by a single TATAA interruption (potentially a mutated TAAAA unit absent from the reference), was found in 55 percent of alleles and exhibited nearly identical repeat lengths. In contrast, alleles composed of pure TAAAA repeats with no interruptions had a median length of 60 base pairs (range: 30–105) (Figure 3F).

In the more ancestrally diverse ONT dataset, the frequency of sequence configurations again varied by superpopulation (Supplementary Figure 1). Among East and South Asian individuals, 42 percent and 44 percent of alleles matched the hg38 reference configuration that lacked the TATAA interruption. In comparison, 55 percent of alleles in the 1,027 All of Us HiFi samples and 58 percent of alleles among individuals of African ancestry in the ONT cohort contained the TATAA interruption. Expanded alleles were only found in configurations lacking the TATAA interruption. However, these expanded alleles were rare, occurring in just 1.29 percent of samples, though the rarity of expanded alleles makes it unclear whether this trend is generalizable.

#### Identification of non-canonical repeat motifs and chimeric repeat alleles

Extending this analysis to all 28,446 loci, 17% of loci had more than one common sequence haplotype seen in at least 5% of alleles (Figure 4A). That is, variation can often extend beyond simple copy number differences and involve distinct configurations of motif and interruption commonly at these loci.

**Figure 4.**
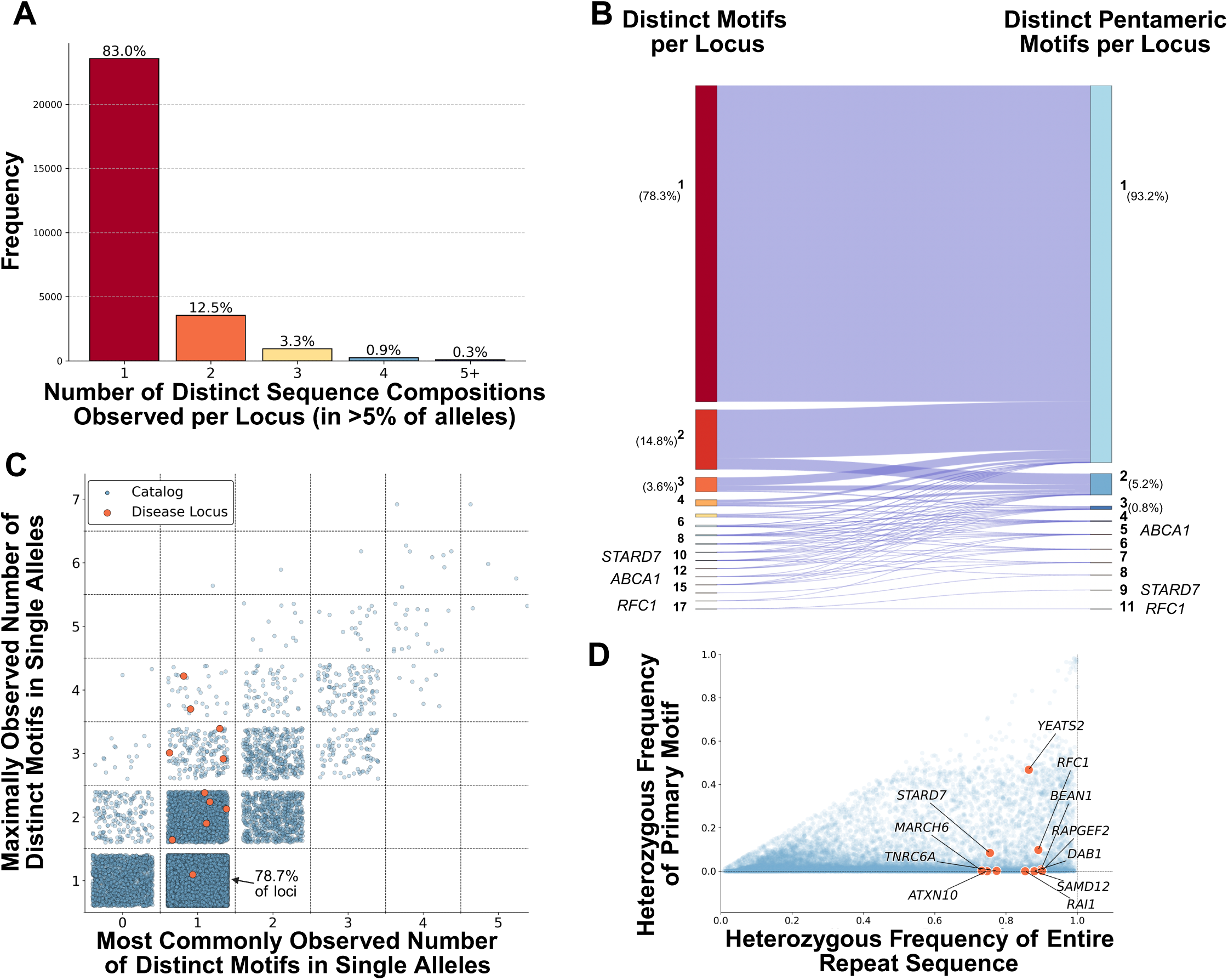
Sequence and motif heterogeneity at pentanucleotide tandem repeat loci. A) Bar plot showing the number of distinct sequence compositions (i.e., unique arrangements of motifs and interruptions) per locus, considering only those observed in at least 5% of alleles at a given locus. Across 28,446 loci in 1,027 genomes, the majority (83%) exhibit a single dominant sequence configuration, while loci with two or more configurations are increasingly rare. Percentages are annotated above each bar. B) Sankey diagram illustrating the relationship between the total number of distinct motifs observed per locus (left) and the number of those motifs that are pentameric motifs (right). Most loci contain a single motif type (78.3% overall, 93.2% when restricted to pentamers), while a subset exhibits motif heterogeneity, with some loci containing up to 17 distinct motifs. Loci in *RFC1, STARD7,* and *ABCA1* are highlighted as examples of extreme motif heterogeneity. C) Scatter plot comparing the most commonly observed number of distinct motifs in single alleles at a locus (x-axis) versus the maximum number of distinct motifs observed in any single allele (y-axis). Most loci (78.7%) contain alleles with only a single motif. Disease-associated loci are enlarged and shown in red. D) Scatter plot of the heterozygosity of the primary motif (y-axis) to the heterozygosity of the entire repeat sequence (x-axis) across 28,446 loci. The primary motif is defined as the motif contributing the most base pairs at a locus. Each point represents a locus, with disease-associated loci shown in red and annotated by gene.

Across the genome, 6,141 (21.7%) of repeat loci contained more than one distinct repeating motif across all alleles (Figure 4B). At loci with multiple distinct motif segments, the most common non-canonical motif was typically either present in fewer than 10% of alleles or in more than 90% at the locus (Supplementary Figure 5). Overall, 21.4% of loci where alleles typically contained a single motif, contained alleles with multiple distinct motifs segments (Figure 4C). Additionally, 10.4% of loci contained alleles that harbored between two and eight distinct repeat motifs within single alleles.

A defining feature of pathogenic intronic pentanucleotide repeat expansions identified to date is the presence of non-canonical motifs within expanded alleles, often resulting in multi-segment repeats that deviate from the reference motif. At known disease-associated loci, the canonical repeat motif is typically presumed benign (e.g., AAAAT, AAAAG, and ATTCT), yet pathogenic alleles contain alternative motifs. We defined a distinct motif as a repeating unit that repeats at least four times consecutively for a minimum of 15 base pairs. In this cohort of only 1,027 samples, the *RFC1* locus, associated with CANVAS, exhibited the greatest number of distinct motifs, with 17 unique repeat motifs genome-wide (Figure 4B). With 11 distinct pentameric motifs, this locus also had the most distinct motifs when only considering pentamers. Similarly, the *STARD7* locus, implicated in FAME2, contained 10 distinct repeat motifs across the studied genomes, with 9 distinct pentameric motifs.

The *STARD7* locus had one of the smaller maximum repeat lengths in the disease-associated set (596 base pairs) but displayed a highly diverse array of sequences (Figure 5A). Approximately 2.8% of alleles were chimeric, containing rare, alternative motifs with or without the canonical AAAAT motif. Many of these recurrent base and motif interruptions were consistently positioned across multiple samples, supporting their authenticity. For example, a single nucleotide difference in the flank of *STARD7* alleles nearly perfectly differentiated expanded (AAACT)n alleles from expanded (AAACT)n(AAAAT)m chimeric alleles.

**Figure 5:**
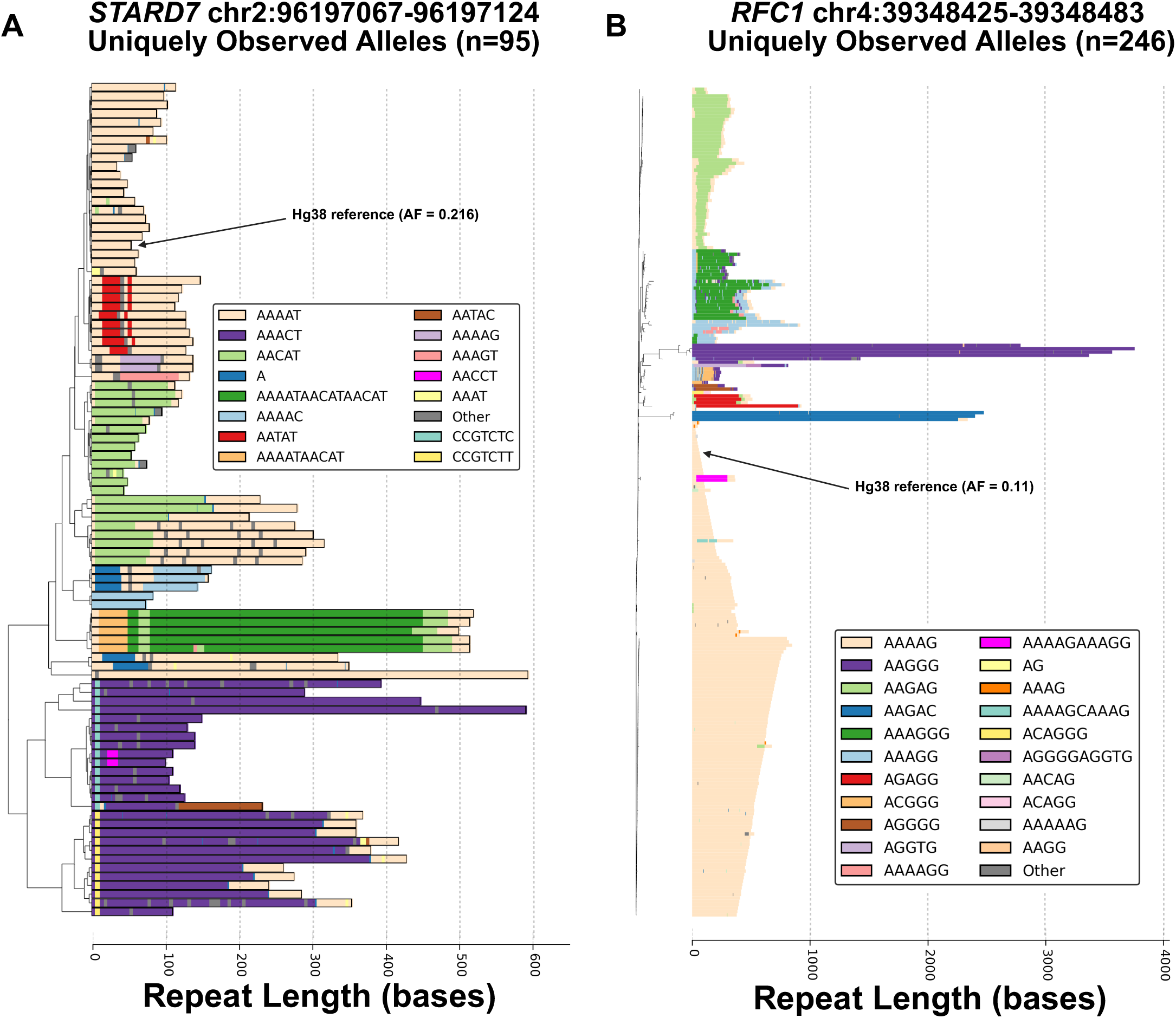
Detailed allelic comparisons of tandem repeats sequences at disease-associated loci. A) Dendrogram of uniquely observed alleles at the *STARD7-*FAME2 locus in 1,027 genomes (N = 95). Each row represents a unique allele, with colored bars indicating repeat motifs or interruptions as specified in the legend. The x-axis represents sequence length in base pairs. Clustering was performed using the Unweighted Pair Group Method with Arithmetic Mean (UPGMA) algorithm, and distances between alleles were computed using a customized edit distance algorithm that accounts for insertion, deletion, substitution, duplication, and contraction as allowable transformations. The hg38 reference allele (AF = 0.216) is annotated. B) Dendrogram of uniquely observed alleles at the *RFC1-*CANVAS locus in 1,023 genomes (N = 246). Each row corresponds to a distinct allele, color-coded by motif composition. The x-axis represents repeat length in base pairs. Clustering was performed using the Neighbor-Joining algorithm, and single-unit interruptions were removed. The hg38 reference allele (AF = 0.11) is indicated.

Further investigation of disease-associated loci revealed substantial sequence heterogeneity. At the *RFC1* locus, 5.57% of alleles contained a non-canonical primary motif. Single alleles contained up to four distinct repeating motifs (Figure 5B). Some alleles also contained decanucleotide motifs such as AAAAGCAAAG, AGGGGAGGTG, and AAAAGAAAGG—features that have not been previously reported or characterized. Additionally, a novel ACAGGG motif was identified in a chimeric allele with the ACAGG motif, illustrating variability in both motif sequence and periodicity. The cohort of 465 ONT genomes identified more unreported motifs in chimeric, expanded alleles such as AAGGGG, AAGAT, AAGACAAGAT (Supplementary Figure 6). This expands the possible motifs associated with the *RFC1* gene though it remains unclear whether these rare repeat structures have any clinical relevance or contribute to disease in a recessive context.

#### Heterozygosity of primary motif

Most known pentanucleotide repeat disorders are inherited in an autosomal dominant manner, with the *RFC1* locus associated with CANVAS being an exception. In this cohort, the disease-associated AAGGG motif at *RFC1* was observed in 1.22% of alleles, prompting investigation of heterozygosity – the presence of different repeat alleles on each chromosome within an individual - across all loci.

Overall, repeat loci displayed a median heterozygosity of 27.1% for total repeat sequence. However, disease-associated loci exhibited significantly higher levels of heterozygosity, ranging from 73.1% to 90.1% (Figure 4D). In contrast, motif-level heterozygosity—defined as cases where different primary motifs occur on each allele (where “primary” refers to the motif with largest base-pair contribution in an allele sequence)—was rare genome-wide. The median motif-level heterozygosity was 0.58%. Eight of the eleven disease-associated loci exhibited motif-level heterozygosity frequencies below 1%. The *ATXN10*, *SAMD12*, *RAI1*, and *DAB1* loci had frequencies of 0%, with all alleles having the canonical motif as the primary motif in this cohort. Exceptions included the *RFC1, STARD7,* and *YEATS2* loci, which exhibited motif heterozygosity rates of 9.8% and 8.4%, 46.8% respectively, revealing significant sequence heterogeneity beyond length polymorphism.

### Evolutionary recency of pentanucleotide tandem repeats

#### Conservation of human reference sequence amongst primates

Nearly 70% of the repeat loci identified in this study map to an Alu element, prompting an investigation into the evolutionary formation of the repeat sequence in primates. The reference sequence at these 28,446 loci for every species included in the UCSC multiZ alignment of 30 mammals, including 27 primates was selected for analysis. Figure 6A illustrates the complex evolutionary trajectory of the SCA31-associated *ATXN10* locus, where repeat decomposition and multiple sequence alignment highlight that the currently associated ATTCT motif likely emerged in the common ancestor of humans and apes.

**Figure 6.**
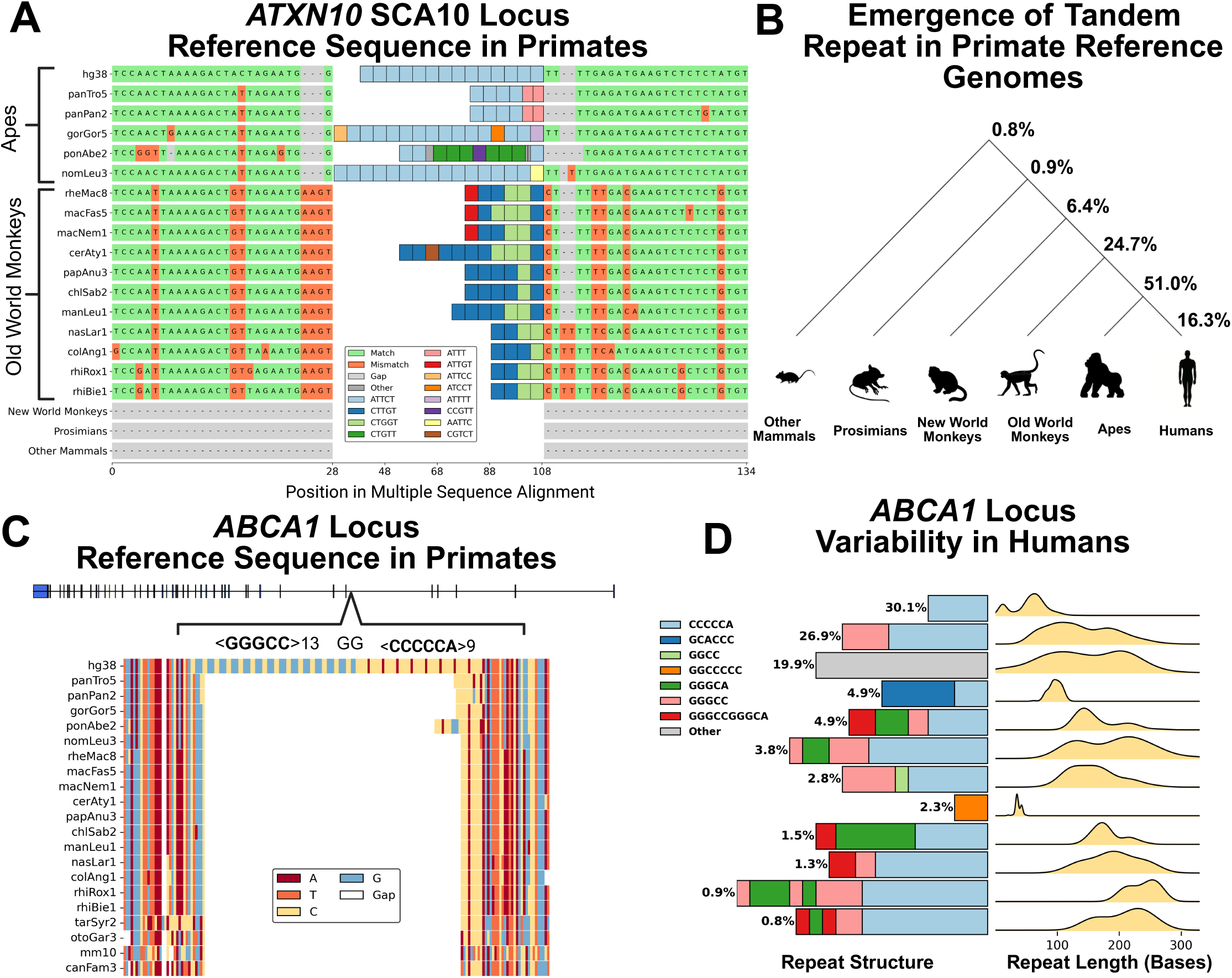
Conservation and emergence of pentanucleotide repeats in primate reference genomes. A) Multiple sequence alignment and tandem repeat decomposition of the *ATXN10* SCA10 locus based on the UCSC multiZ alignment of 30 mammals (27 primates). The flanking sequences were aligned using Muscle version 5.1. Colored bars represent distinct repeat motifs (see legend), with major primate clades bracketed. Matches, mismatches, and alignment gaps are color-coded, and sequence length is shown on the x-axis. B) Schematic illustrating the phylogenetic emergence of 28,446 loci in primates. The earliest genome that contained four uninterrupted repeat units matching the hg38 motif was used to define the group from which the locus emerged. Percentages above each split indicate the proportion of these loci appearing at each phylogenetic node. C) Gene structure of *ABCA1*, highlighting the intron 5 location of the tandem repeat locus. The entire sequence was aligned using Muscle version 5.1, with colors representing distinct nucleotides and white blocks denoting alignment gaps. The sequence and structure of the decomposed repeat in hg38 is written above the sequence. D) Diagram depicting the structure and length distribution of alleles at the *ABCA1* locus (*chr9:104862606-104862843*). Left: A horizontal stacked bar plot representing the motif composition of *ABCA1* repeat alleles. Each color corresponds to a distinct repeating motif, as indicated in the legend. The length of each bar is proportional to the median number of repeat units observed for alleles with that structure. The percentage frequency of each repeat composition in the dataset is annotated next to its respective bar. Right: Kernel density estimation (KDE) plots illustrating the length distribution (in base pairs) of decomposed repeats for each structural composition.

Using ATXN10 as a guide, we extended this analysis to all 28,446 loci to determine in which primate group the human repeat motif first fully emerged. Using a simplified criterion for repeat existence—requiring at least four consecutive units matching the most common human repeat motif— 51% of loci were found to have first fully emerged in an ape lineage, while 16.3% showed no species with a matching four consecutive units (Figure 6B). To further validate these findings, we leveraged the *mPanTro3* diploid chimpanzee genome assembly from the Telomere-To-Telomere (T2T) consortium, aligning it against the hg38 reference genome to assess the 16.3% of loci that appeared human-specific. This updated chimpanzee assembly corrected 233 loci, refining our estimate to 15.4% of loci remaining specific to the human reference genome. Among the known disease-associated loci, eight first emerged in an ape species, and three in Old-World monkeys. The *RAPGEF2-FAME7* repeat locus—the only one not mapping to an Alu element—lacked four uninterrupted TTTTA repeat units until the gibbon reference sequence (Supplementary Figure 7). This suggests that, despite being in a more ancient genomic region, the tandem repeat itself may represent a more recent evolutionary event.

#### Identification of recently evolved and highly polymorphic loci

Integrating these evolutionary findings with human population-level data highlighted loci that are both evolutionarily recent and highly unstable in humans. One example of a locus that was highlighted as ‘Human Specific’ in this analysis mapped to intron 5 of the *ABCA1* gene. This repeat locus is nested inside a MER5A transposable element less than 1kb away from the nearest exon and contains pentanucleotide CCGGG and hexanucleotide ACCCCC repeat units in the hg38 reference sequence (Figure 6C). Despite the MER5A transposable element existing in the reference sequence of apes and old-world monkeys, the human-associated tandem repeat was not present in any other reference genome. This locus overlaps with an H3K27Ac mark in GM12878 cells (lymphoblastoid) based on ENCODE ChIP-seq data and is transcribed on every *ABCA1* transcript. In the human population, this locus possessed exceptional sequence heterogeneity and complexity: there were 1,210 distinct decomposed repeats at this locus, and 13 distinct motifs identified across 2,050 alleles (Supplementary Figure 8). Approximately 5.4% of alleles were composed of only two copies of the ACCCCC motif. When removing static interruptions, we identified 9 sequence configurations of variable motif copy numbers and frequency (Figure 6D). Despite only having an expansion score of 2.66 (maximum 329, median 90 base pairs), this region was among the most sequence diverse repeats in the entire genome. This locus exemplifies a broader class of highly variable, evolutionarily recent repeats that remain underexplored in studies of human health and disease.

## DISCUSSION

### Advancing Genomic Analysis of Tandem Repeats

Tandem repeats are often oversimplified as a motif unit and count, an approach that overlooks the complexity of repeat configurations and veils potential phenotypic implications. This study advances the field by showcasing detailed and comprehensive characterizations of tandem repeat variation.

Customizable algorithms were developed for repeat decomposition, comparison, and visualization. These methods uncovered recurrent sequence variants and configurations that would likely remain hidden using conventional approaches. By visualizing the sequence composition of alleles and deriving common sequence configurations, we discovered that distinct sequence compositions dominate distinct length ranges at loci. For loci with multiple rare sequence configurations, such as *STARD7* and *RFC1*, phylogenetic tree-based profiling using a customizable edit distance algorithm— including motif duplication and contraction—provided clearer insights. Based on existing examples, pathogenic repeat alleles likely exhibit higher edit distances from the reference sequence, offering a potential framework for automated prioritization of candidate loci and outlier genotypes. Importantly, our approach enables automated solutions for repeat characterization that maintains high-resolution capture of repeat motif and interruptions, like we showed for *SAMD12*, processes previously requiring intensive manual inspection to detect such subtle differences. These methods allow for large-scale screening to identify clusters of variation and facilitate detailed allelic comparisons. One such application would be clarifying different allele frequencies between populations when considering the sequence content of an allele. Multiple sequence alignment demonstrated the importance of incorporating flanking sequence variation into analyses. This approach could help identify imputable variants for sequence haplotypes, such as those we identified at *STARD7*, or biologically relevant flanking variations, as exemplified by the *FGF14* repeat locus associated with SCA27B [32].

### Current limitations and future goals

This dataset of 1,027 genomes analyzed here represents one of the largest collections of HiFi data analyzed to date; however, it should not be considered exhaustive. The catalog of motifs and genotypes presented here reflects a conservative lower bound. As additional long-read datasets become available, the number of distinct motifs and sequence configurations will expand.

The HiFi dataset consists of self-identified Black or African American individuals. Some studies note that African genomes contain excess tandem repeat variation, which could translate into enrichment of diversity in this cohort [33]. Expanding the dataset to encompass individuals from diverse ancestries is essential for capturing the full spectrum of tandem repeat variation, including ancestry-specific patterns [7, 34]. At the *SAMD12*–FAME1 locus, we did not observe any large AAAAT repeat expansions, with the maximum observed length reaching only ∼30 repeat units. This contrasts with previous studies reporting expansions exceeding 80 units in 3.4–5.9% of Chinese and Japanese control populations [23, 24]. Reported cases of FAME1 are primarily concentrated in Asian populations, including Japan, China, India, Sri Lanka, and Thailand [35]. In this cohort, the *hg38* reference repeat configuration was present in only 7% of alleles. Instead, two alternative configurations, including a TATAA-interrupted haplotype, were more frequently observed. The smaller but more ancestrally diverse cohort of 465 ONT samples analyzed hinted that alleles with the TATAA interruption are more stable in length, whereas those without it can expand to thousands of base pairs long. Further, the distribution of repeat structures differed significantly by superpopulation with the largest alleles being in east and south Asian superpopulations. Although the mechanisms underlying expansion and pathogenicity of FAME1 are not fully understood, population-specific differences in repeat architecture with different levels of stability may contribute to ancestry-associated variation in disease prevalence.

Phylogenetic analysis of reference genomes from 27 primates employed a heuristic definition of repeat existence to enable automated processing. We selected a threshold of four uninterrupted repeat units, consistent with our criteria used for selecting human loci (see Methods). Multiple genomes per species group (e.g., Prosimians, New World monkeys, Old World monkeys, and Apes) enhanced confidence in the evolutionary trajectory of certain loci. However, this analysis was limited to reference genomes (one per species), therefore did not capture intraspecies variation. As a result, we conservatively defined the emergence of repeats at group levels (e.g., Apes or Old-World Monkeys) rather than pinpointing specific species. Future studies incorporating a greater number of primate genomes per species and modern long-read assemblies will enhance our understanding of the evolutionary origins of tandem repeats. Larger datasets would also allow for cross-species comparisons of repeat length and copy number, a strategy that could reveal loci uniquely expanded in humans [36, 37]. The loci analyzed in this study were all present in the human reference genome. However, alternate strategies could identify tandem repeat loci present in non-human species that have contracted or been deleted from the human genome.

Pentanucleotides were prioritized for this analysis because they represent a subclass of pathogenic repeats most relevantly suited for investigating sequence composition in relation to disease. Extending this methodology to other classes of short tandem repeats will elucidate genome-wide sequence heterogeneity and variability. Tangentially, this work necessitates improved methods to characterize mini-satellites (motif lengths >10 bp) and macrosatellites (motif lengths in the hundreds or thousands) in the human genome. For larger or more complex repeat units, requiring an exact decomposition of repeat sequence can be prone to errors and may require alternative analytical frameworks [38, 39].

### Insights into genome-wide variability of pentanucleotide repeats

Long-read sequencing has revolutionized the study of large and complex tandem repeat alleles, enabling sequence-level resolution that was previously unattainable. Our results reveal a panoply of length and sequence heterogeneity at pentanucleotide loci, highlighting their evolutionary recency and unique features.

Despite their established connection to disease, many pathogenic repeat loci have only been thoroughly validated and characterized within their original discovery cohorts. Larger-scale studies using short-read sequencing have been unable to resolve sequence composition or fully elucidate expanded alleles. Therefore, we examined variation at known disease-associated loci and extended the analysis to comparable loci across the genome. Disease-associated loci exhibited pronounced motif and length heterogeneity in the population, a contrast to traditional interpretations of genetic constraint in disease-causing genes. These loci had heterozygous genotypes in 73% to 90% of samples and had significantly higher expansion scores than other loci. Although the individuals in this dataset were not selected based on disease status, deleterious repeat alleles with incomplete penetrance or recessive inheritance, such as those at *RFC1*, remains a possibility. Additionally, tandem repeats in compound heterozygous state with other genetic variations could contribute to undiagnosed conditions. Thus, the motif and length frequencies of unstable loci identified in this study provide valuable data for investigating potentially overlooked pathogenic genotypes.

Our analysis clarifies seldomly discussed features of variation at highly polymorphic tandem repeat loci. Non-canonical motifs were present at most disease-associated loci, demonstrating their natural occurrence beyond pathogenic genotypes. Often, these motif changes were present in alleles with multiple distinct repeating units. Sequence configurations with different motifs and structures often occupied distinct length ranges at a given locus, and subtle but recurrent interruptions would often separate otherwise similar alleles. For instance, the major configuration we observed at the *SAMD12*- FAME1 locus in the HiFi cohort occupied the same length ranges as the configuration found in the hg38 reference genome but differed by a single base-pair interruption. These findings underscore the limitations of defining alleles solely by motif and unit count, emphasizing the need for more nuanced databasing approaches.

Our high-resolution approach to repeat decomposition provided insights into the assembly and evolution of repetitive DNA sequences. Previous studies give two hypotheses for the creation of chimeric tandem repeats: mutation of the existing motif or the insertion of new repeat motif. In our results, we see evidence for both mechanisms. It was noticeable that distinct adjacent motifs in alleles often differ by a single base pair. For example, at the RFC1 locus, for instance, a novel ACAGGG motif was observed adjacent to the disease-associated ACAGG motif within an allele. This could reflect a mutation in the current motif followed by duplication of the altered unit. This mechanism also supports the observation that polyadenine sequences in *Alu* elements contribute to the genesis of these repeat loci, as the most common pentameric motifs in the human genome (AAAAC, AAAAT, AAAAG) differ by only a single mutation from a homopolymer adenine sequence. Additionally, at the *RFC1*, *STARD7*, and *YEATS2* loci, we observed decamer and 15-mer motifs, which may represent an initial motif mutation followed by the fusion of units into longer motifs. For the second mechanism: waterfall visualizations and sequence haplotype analysis of many loci demonstrate that specific configurations cluster separately in length, without a spectrum of intermediate alleles that demonstrate the formation of them. This may reflect a single disruptive event creating the formation of a more stably inherited sequence haplotype rather than gradual stepwise mutation. Further, alleles with non-canonical motifs were often rare and presented in the sequence terminus of the largest alleles at a locus. Ultimately, studies examining both inherited and somatic tandem repeat mutations beyond simple copy number changes will provide further insight into these observations.

Pentanucleotide tandem repeats are strongly enriched in proximity to Alu elements (68%) and other transposable elements (16%), despite Alu elements comprising only 11% of the human genome [40]. Phylogenetic analysis of non-human primate reference genomes further suggests that the majority of pentanucleotide repeats first “emerged” in a common ancestor of humans and apes or potentially in genomes more closely related to humans. These analyses identified loci specific to human genomes— variations that may contribute to speciation and the emergence of human-specific traits. The tandem repeat locus we highlight in *ABCA1* is ultra-polymorphic in the population but may only exist in humans. The *ABCA1* gene plays a key role in regulating high density lipoprotein (HDL), cholesterol metabolism, and cardiovascular disease [41]. This repeat locus is located close to an exon (<1 kb) and overlaps with an H3K27Ac mark and a distal enhancer-like signature. Despite its proximity to these functional regions, studies regarding *ABCA1* may be overlook this form of variation due to its location outside protein-coding regions.

To conclude, we have identified the loci in the human genome that are prone to instability, yet the mechanisms underlying this phenomenon remain unclear. Disease-associated loci are among the most polymorphic in the genome, raising the question of whether their instability predisposes them to disease simply because their dynamic nature increases the likelihood of disruptive events, or if specific biological mechanisms actively drive their propensity for instability, with disease as an unintended consequence. Similarly, we pinpoint repeats that have evolved more recently in humans, yet their phenotypic impact and native functions, if any, is an ongoing question. Investigating these aspects may yield critical insights into the selective pressures and functional roles that shaped these sequences. Particularly, studying the extreme cases we identified offer an opportunity to illuminate the biological triggers of repeat expansion or creation. Understanding the causes of this instability may hold the key to therapeutic development, as somatic repeat instability is increasingly linked to disease pathogenesis [42, 43]. Ultimately, the broader significance of tandem repeat variability, including its contributions to human uniqueness, genome evolution, and susceptibility to disease, represents a rich and underrecognized area of genomic discovery.

## METHODS

### Data processing and tandem repeat calling

A cohort of 1,027 individuals were selected for long-read sequencing as part of the All of Us Long Reads Working Group. All of these individuals self-identified as Black or African American. Blood samples had been collected for each of these individuals and stored in accordance with the All of Us Research Project protocols. High molecular weight DNA was extracted and sent to the HudsonAlpha Institute for Biotechnology (Huntsville, AL) for sequencing. Samples were sequenced to an average coverage of 8x using Sequel IIe machines (Pacific Biosciences).

Genomes were aligned to the GRCh38 reference genome assembly using pbmm2 version 1.7.0. Tandem repeats were genotyped using TRGT version 0.3.3 with the Adotto catalog version 1.0 (https://zenodo.org/records/7521434). Repeat loci were padded by a minimum of 25bp on each side and loci longer than 1,000bp in the GRCh38 reference genome removed to accommodate a limitation of the version of TRGT used.

### Data processing and tandem repeat calling for 465 Oxford Nanopore Technology (ONT) samples

We analyzed a cohort of 465 publicly available whole-genome sequences generated by the 1000 Genomes Project ONT Sequencing Consortium [44]. For select loci discussed in this paper, STRdust (https://github.com/wdecoster/STRdust) was used to generate a consensus sequences for each sample [45]. The ancestry superpopulation given in the International Genome Sample Resource (IGSR) data portal was considered for each sample.

### Selection of Pentanucleotide Repeat Loci

The longest pure segment (LPS) was computed for 1,757,271 catalogued loci in 1,027 PacBio HiFi samples. Loci that at least 100 alleles with a pentanucleotide LPS motif were selected if they had a median length of at least four units or had a minimum difference between the median and maximum LPS of 20 base pairs. Less than 0.3% of loci with a median LPS less than 20 base pairs had a range from median to maximum LPS of at least 4 units. This totaled to 28,446 pentanucleotide repeat loci. Telomeric and centromeric regions were not considered. For this analysis we considered motifs analogous by any process of reduction, shifting, or reverse complementation and labeled in minimal alphabetical order. RepeatMasker [46] annotations for the hg38 reference genome were computed using bedtools [47]; for loci that intersected with more than one element, the element that had a higher bp overlap was selected.

### Decomposition of Tandem Repeat Alleles

Observing tandem repeat sequence interruptions and variants in multiple samples gives increased assurance of their authenticity. However, commonly used tandem repeat decomposition tools implement an approximate tandem repeat genotype with a mismatch percentage [48, 49]. Therefore, we devised an algorithm that will attempt to compute an exact tandem repeat decomposition from a string. This algorithm is a heuristic algorithm we have named RepeatXuTRACTR (repeat extractor) that segments a string using a library of repeating kmers found in the string. For alleles with multiple repeat decompositions, the decomposition with the longest pure segment was selected for analysis. An example of the input and output of our algorithm is shown below.

Input: GCAACCACACCCGGCCTAGAGAAGCGTTCAGTTCAGTTCAGTTTAGTTTAGTTTAGTTTAGTTTA GGTTAGGTTAGGTTAGGTTAGGTTAGGTTAGGTTAGGTTAGGTTAGGTTAGGTTAGTTTAGTTTAGGTTTAGTTTAGTTTAGTTTTAGTTTTTGAGACGGAGTCTCGCACTTTCA

Output: <AGTTC>2<AGTTT>5<AGGTT>11<AGTTT>2<AGGTT>1<T>1<AGTTT>2<AGTTTT>2

### Derivation of common sequence haplotypes

The output of our string decomposition algorithm was grouped by structure, defined by the order of motif units and interruptions. This method assumes that repeat motif duplication or contraction precedes substitution and or indel mutations. Each haplotype was represented as a string formatted as <MotifA>min-max<MotifB>min-max where min and max indicate the minimum and maximum number of units for that motif for alleles with that specific structure. For non-variable interruptions or instances where these values were equal, a single value was used instead.

### Repeat Expansion Scores

For each repeat, an expansion score was calculated as [(Maximum repeat length – median repeat length) / median repeat length] [50, 51]. This metric was used to capture positive skew and outliers at a locus, a feature of some tandem repeat loci associated with repeat expansion disorders.

### Creation of a phylogenetic tree for tandem repeat sequences

A pairwise distance matrix was computed using an extension of the Levenshtein distance that incorporates duplication and contraction of motifs as possible transformations in addition to insertion, deletion, substitution. Various implementations of this logic have been priorly utilized for comparing minisatellites [52–54]. We customized this algorithm that we call XuACTDist (Exact Distance) to use motif allele frequency in our cohort when considering edit costs, as well as motif sequence when calculating motif substitution costs. For instance, rare motifs had a ”higher cost” to insert/delete at a locus, and an AAAAG/ACGGG substitution would have a “higher cost” than an AAAAG/AAAGG substitution. This algorithm was intended to prioritize repeat sequence content (motifs and interruptions) over repeat length when comparing sequences, so duplication and contraction costs were designed smaller than insertion, duplication, and substitution. For a string s length m and string t length n, the algorithm uses O(m*n) space and O(m*n) time complexity. If using allele frequency and motif information, an optional O(k^2^) space complexity was used for a cost dictionary where k is number of distinct motifs.

### Conservation and Alignment of Tandem Repeats amongst Primates

The Cons 30 Primates track in the UCSC genome browser was used to identify genomic coordinates of homologous tandem repeat loci to the hg38 reference genome in 30 species (27 primates). However, the multiz30way alignment is specific for the hg38 reference sequence, and in certain instances will remove sequence elements absent from hg38 but present in multiple other species. Therefore, the reference sequence at these loci for each species was extracted with bedtools getfasta [47] and decomposed using our algorithm. Muscle multiple sequence alignment version 5.1 [55] was used to align the flanks of tandem repeats, and the repeat sequence was visualized in between.

