## Supplemental Figures for "The Genetic and Evolutionary Landscape of Pentanucleotide Tandem Repeats in Human Genomes"

### Slide 1
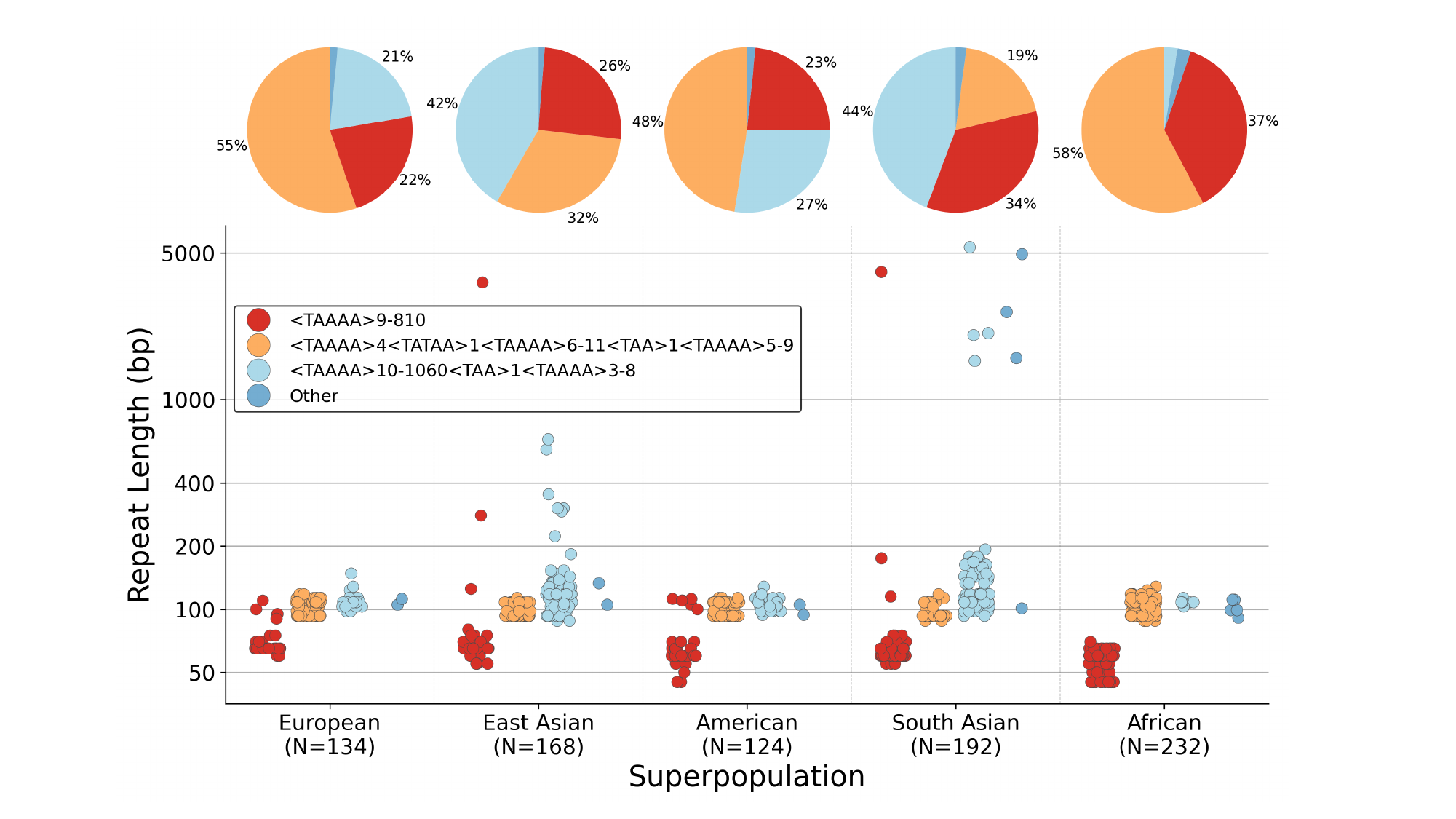

### Slide 2
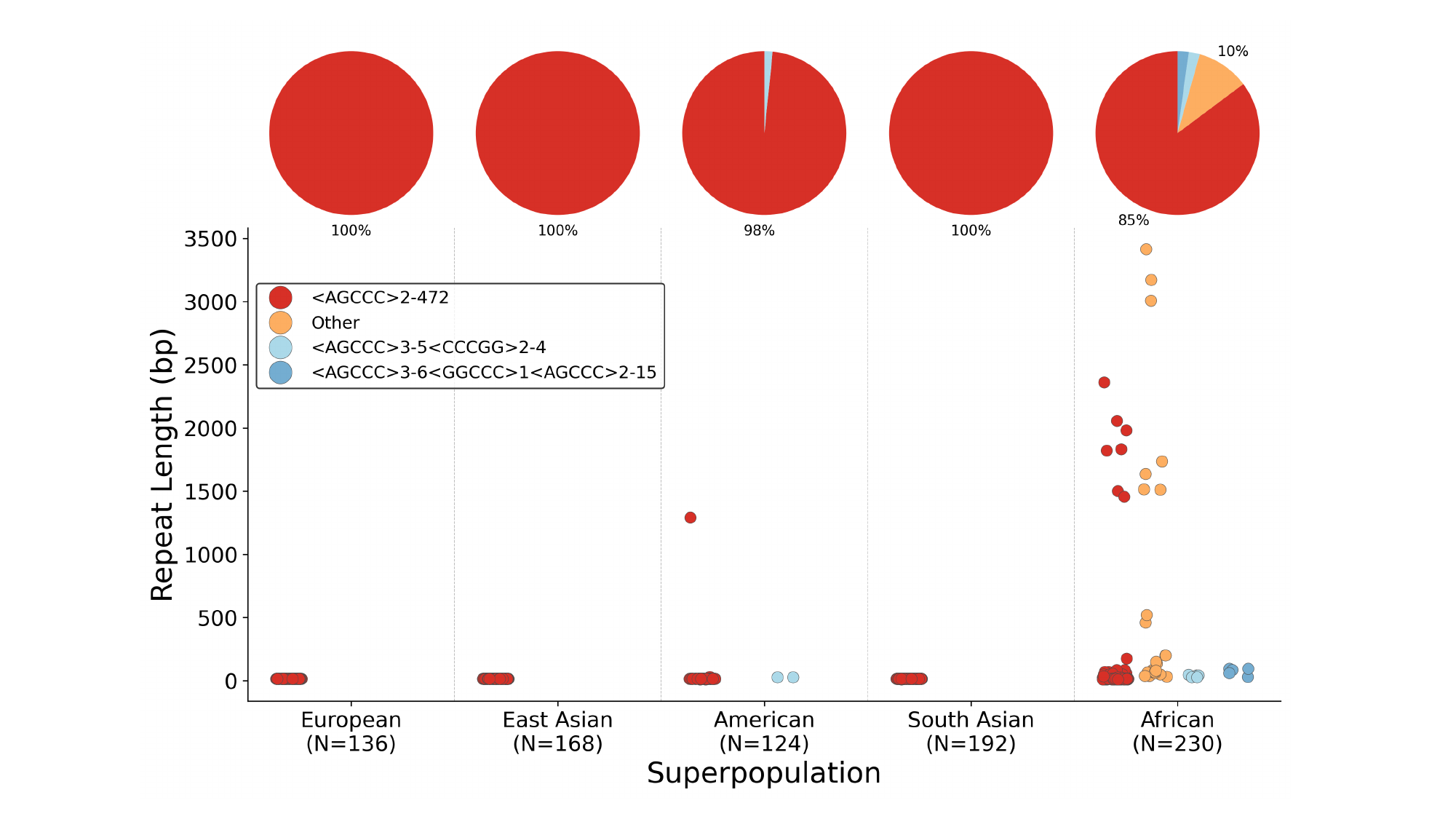

### Slide 3
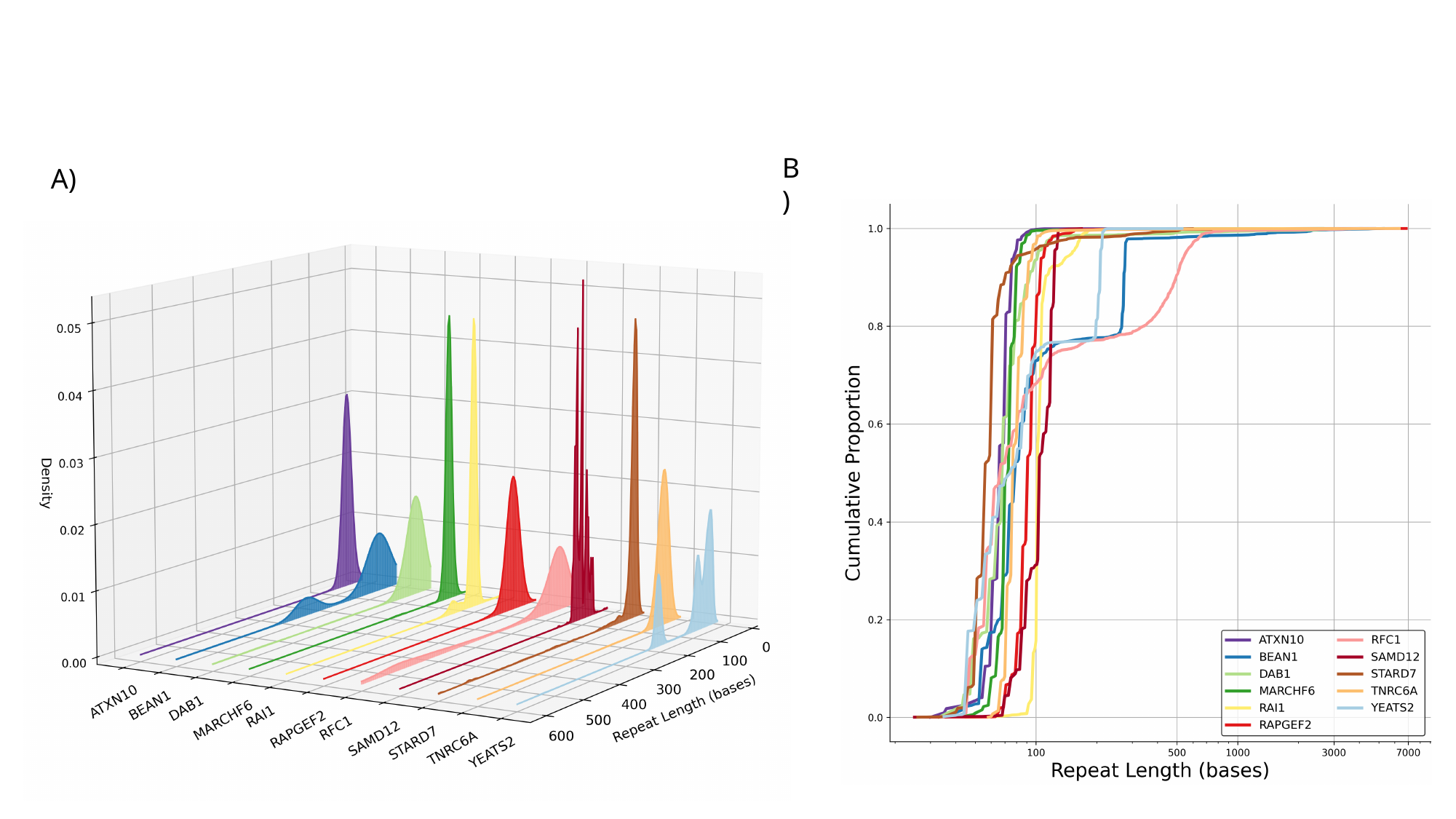

B)
A)

### Slide 4
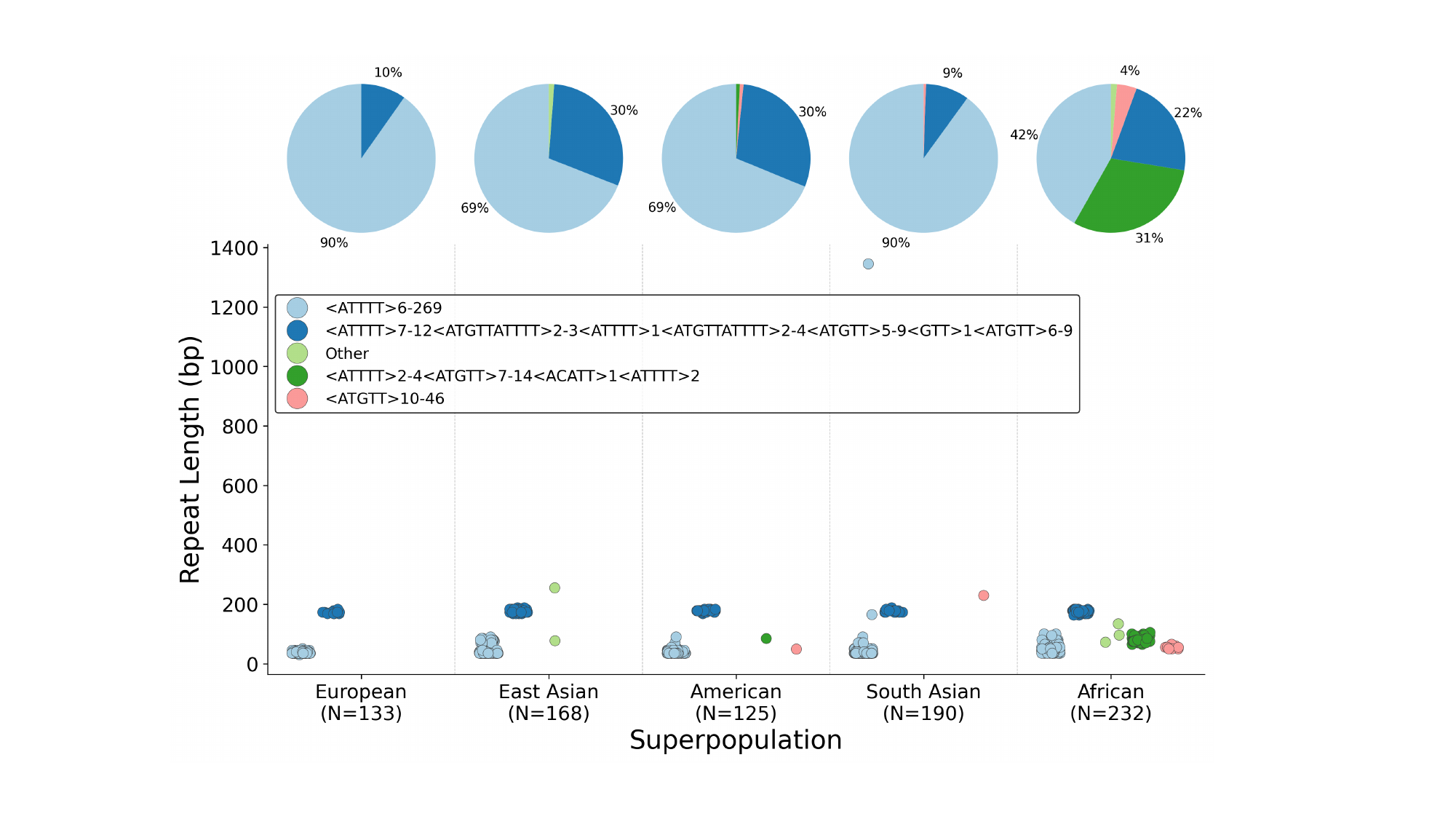

### Slide 5
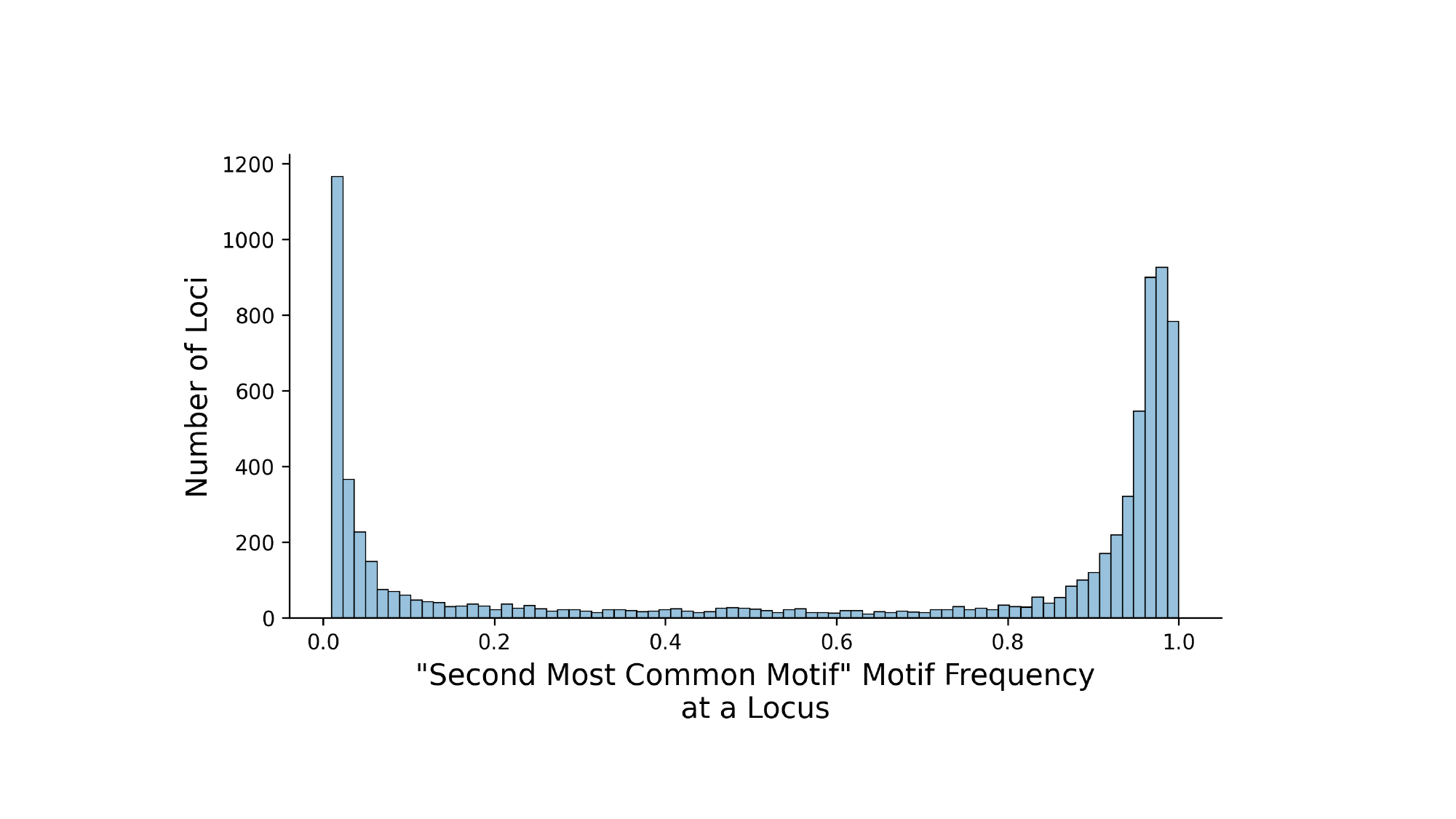

### Slide 6
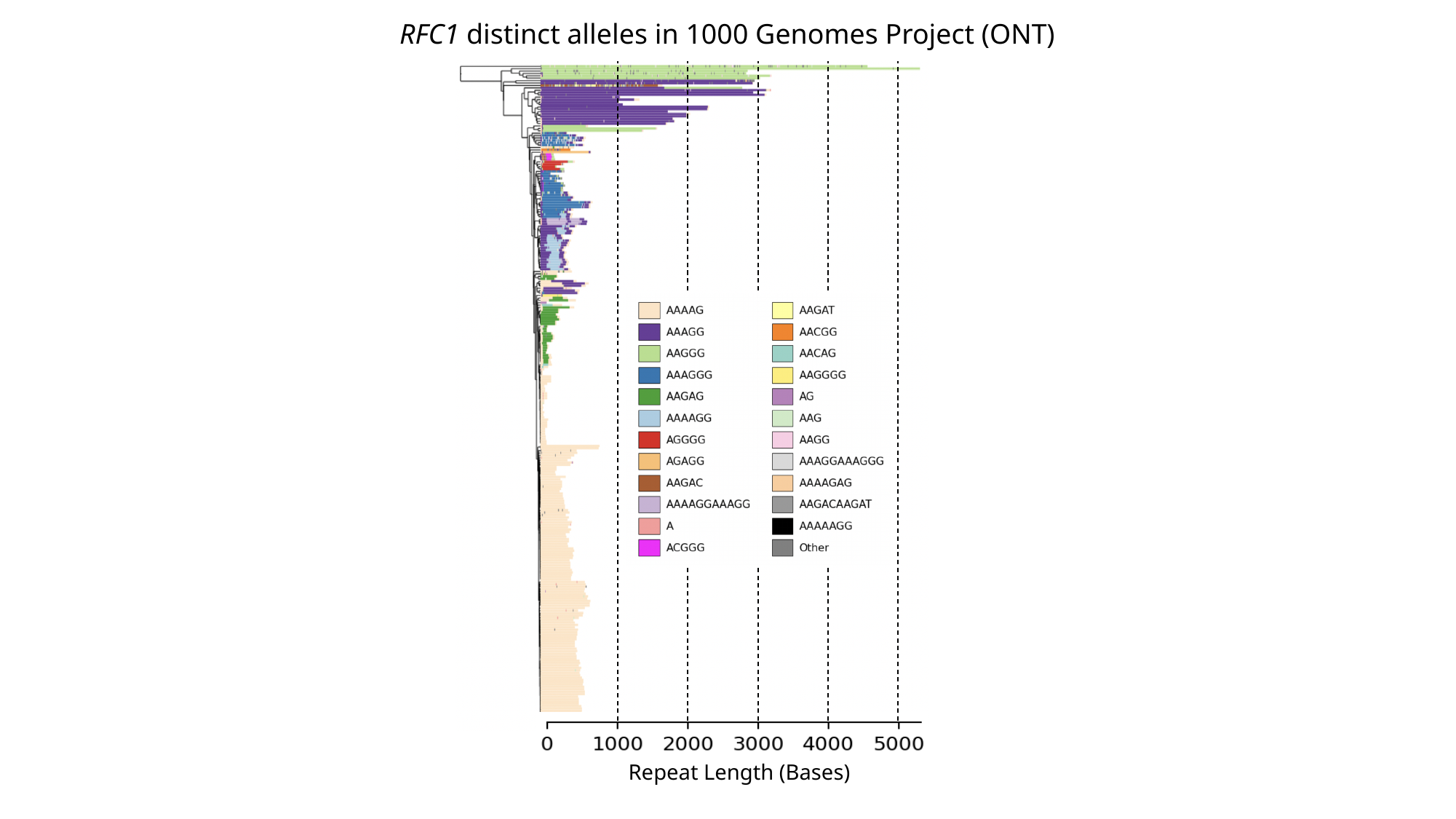

RFC1 distinct alleles in 1000 Genomes Project (ONT)
Repeat Length (Bases)

### Slide 7
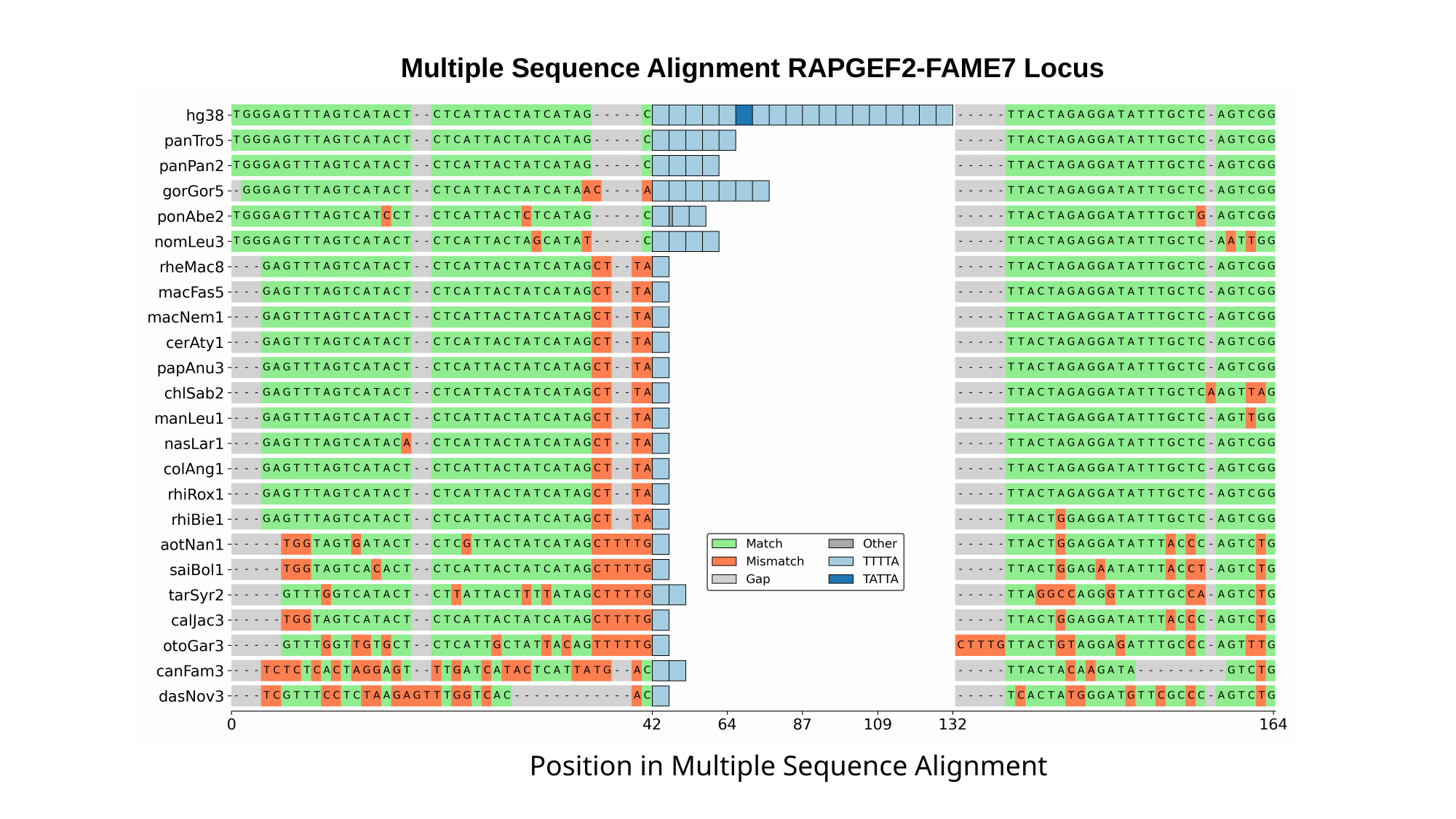

Multiple Sequence Alignment RAPGEF2-FAME7 Locus
Position in Multiple Sequence Alignment

### Slide 8
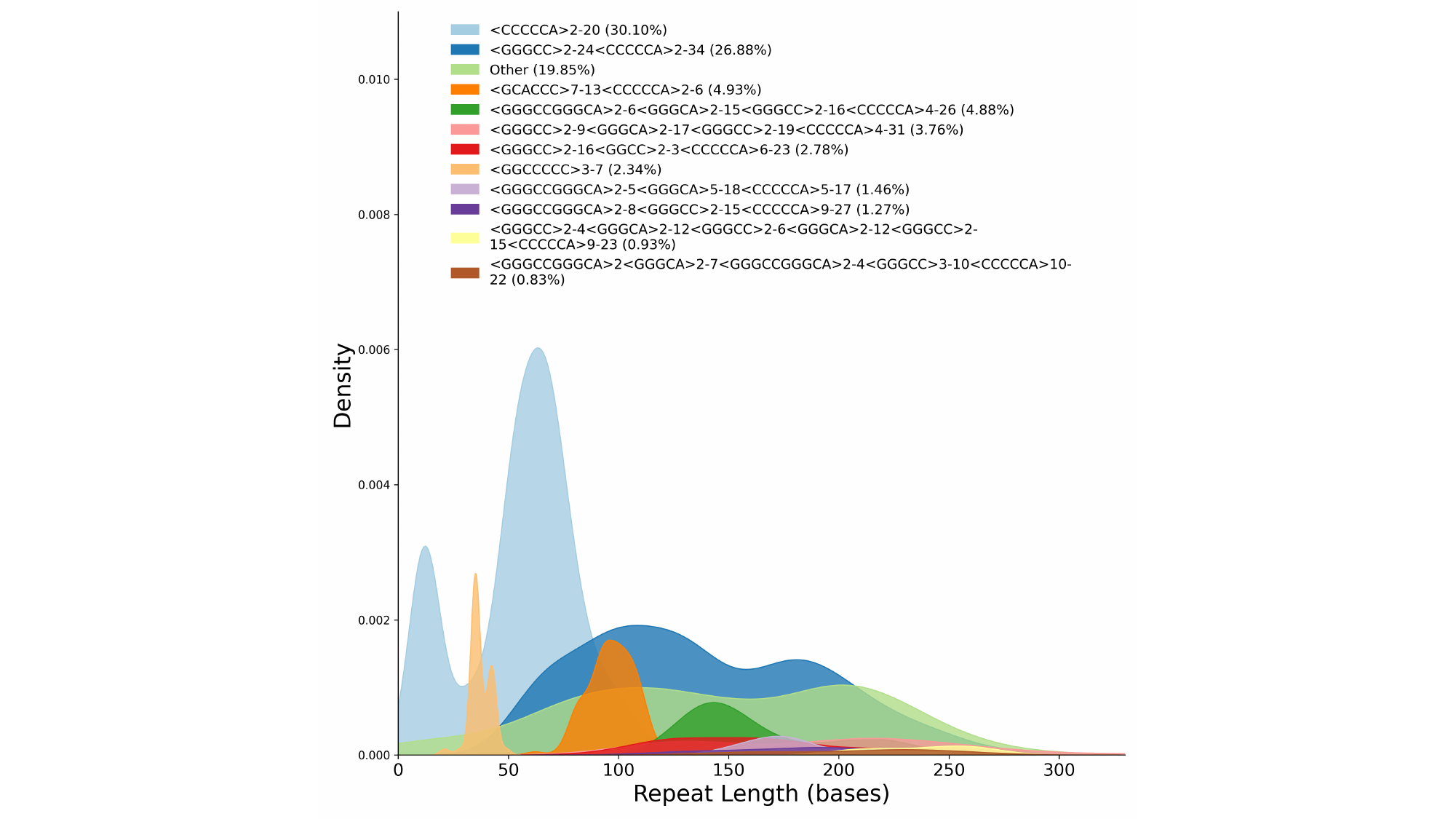
